## Supplemental File for "ThermoMaze: A behavioral paradigm for readout of immobility-related brain events"

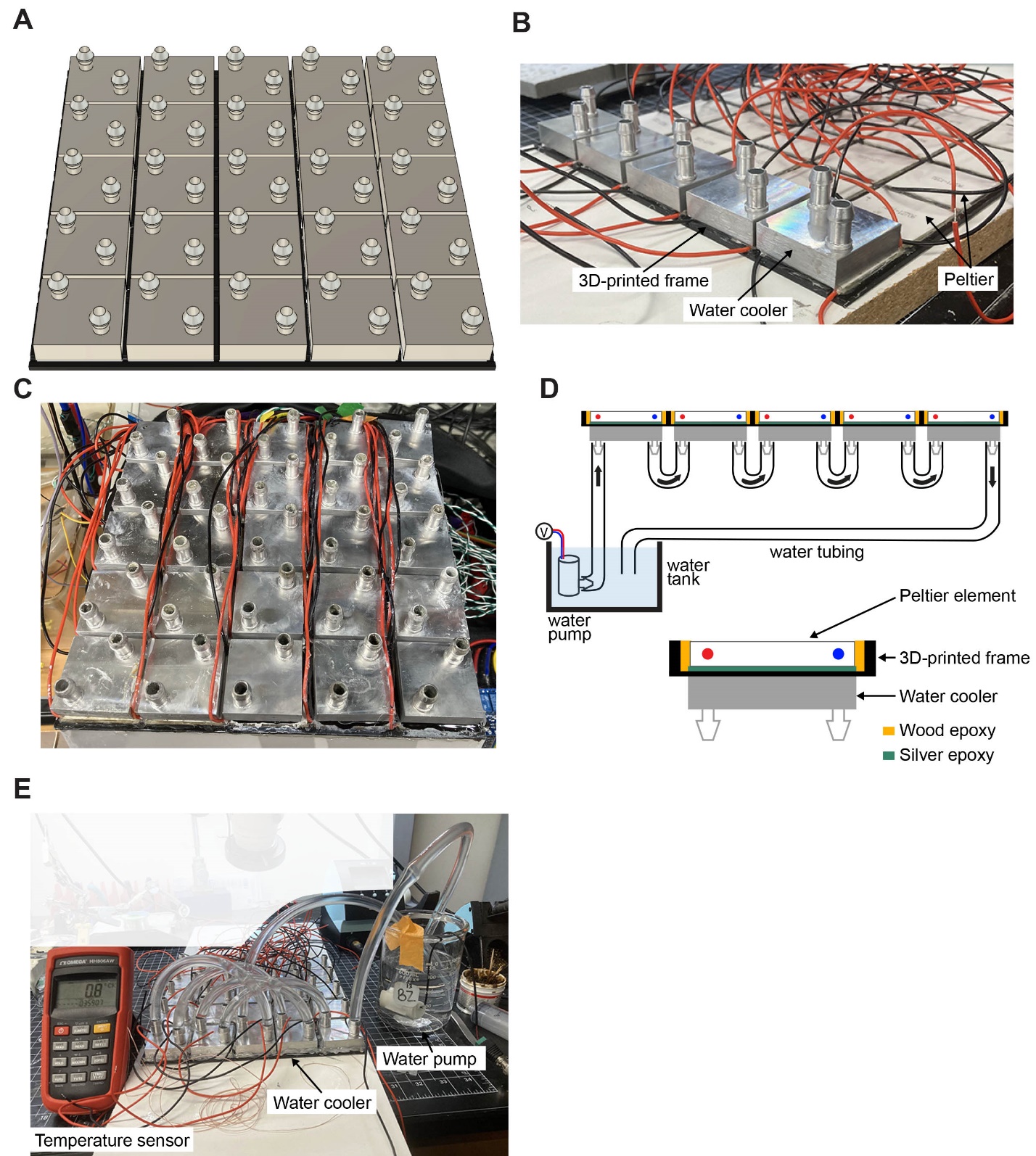

**Supplementary Figure 1. Control of heating and cooling of the surface of ThermoMaze. A)** Schematic of water coolers (each Peltier element has its own water cooler, n = 25). **B)** Photograph of ThermoMaze with all Peltier elements attached to a 3D-printed frame (bottom view). One row of water coolers (n=5) is also attached to Peltier elements. **C)** Photograph of the bottom view of the ThermoMaze showing 25 water coolers without tubing attached. **D)** Schematic of water circulation system. **E)** Ice-cold water circulating through the water tubes and between 5 water coolers and Peltier elements (turned off) can passively reduce the surface of the Peltier element to 0.8 °C. The temperature is measured by a K-type thermocouple attached to the surface of the last Peltier element in a row.

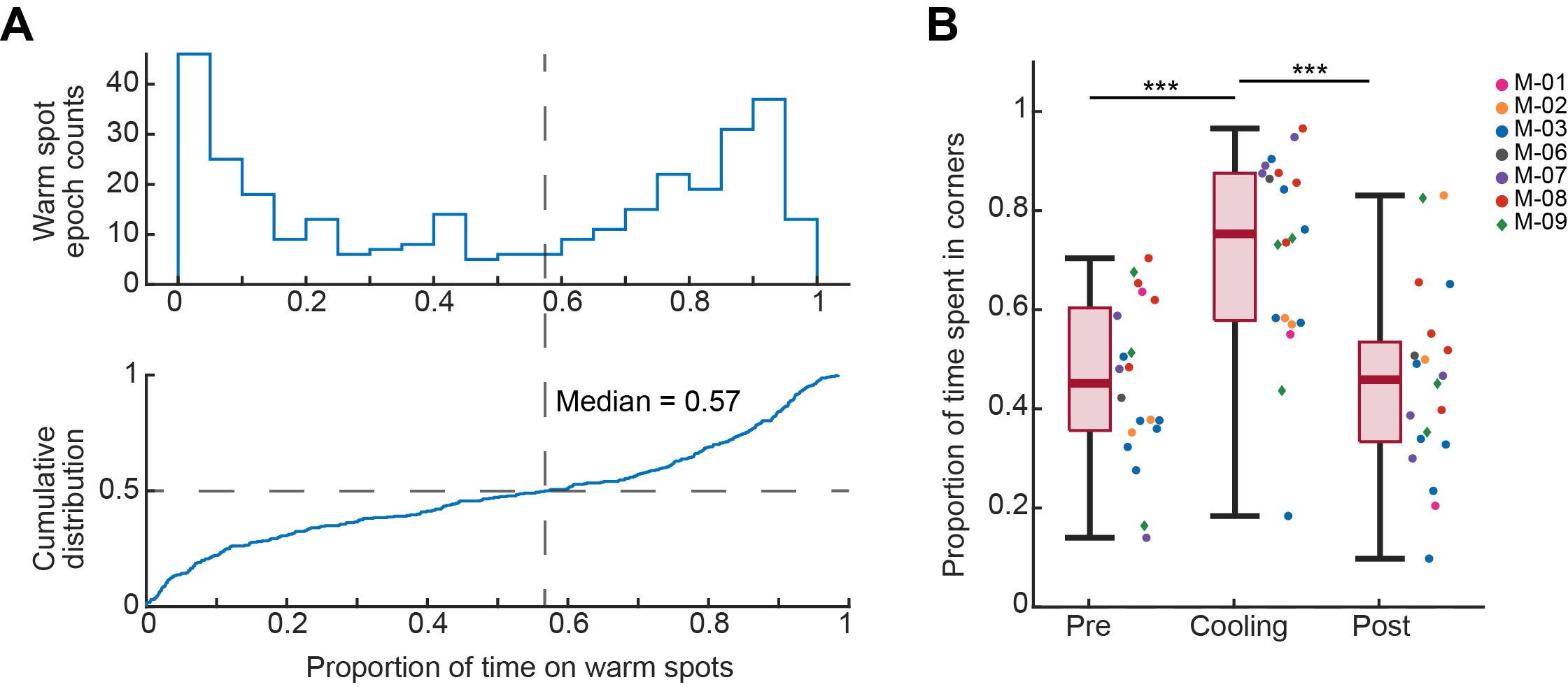

**Supplementary Figure 2.** **Animals learned to track and stay immobile on hidden warm spots in the ThermoMaze.** **A)** Top: histogram of proportion of time spent on the warm spot during each warm spot epoch when it was providing heat. 0 indicates that the animal did not occupy the warm spot when it was turned on, and 1 indicates that the animal was staying on the warm spot for the entire warm spot epoch. Bottom: cumulative distribution of the proportion of time animal spent on the warm spot during a warm spot epoch (median = 0.57; in other word, median = 2.85 minute per 5-minute warm spot transition epoch). Therefore, in over 50% percent of the warm spot epochs, mice found and stayed on the warm spot for over 57% of the time (n = 20 sessions in n = 7 animals). **B)** Box plot of the proportion of time that the animal spent in any of the four warm spot corners in the ThermoMaze. Median, Kruskal–Wallis test: H = 19.69, d.f. = 2, p = 5.29*10^-5^. The proportion of time spent in corners in pre and post and significantly different from cool (Pre vs. Cooling: p = 0.0004; Cooling vs. Post: p = 0.0003), while that of pre and post are not significantly different (Pre vs. Post: p = 0.9996). Dots (females) and diamonds (males) between the boxes represent the individual sessions and the same color represents sessions from the same animal.

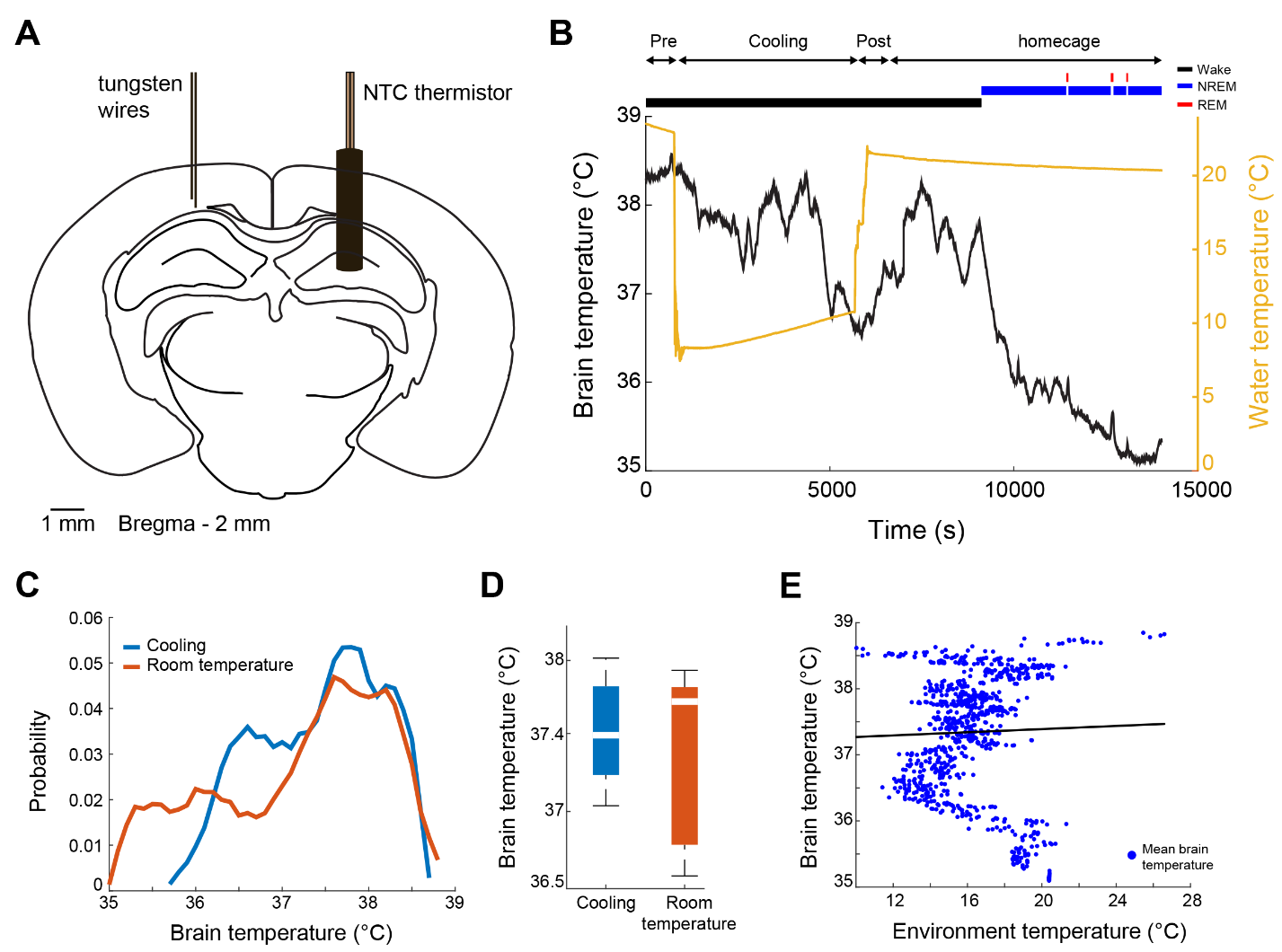

**Supplementary Figure 3. Brain temperature is not affected by cooling of the ThermoMaze. A)** Schematic of implantation of the thermistor. Mice were implanted and tungsten recording wires. **B)** Brain temperature variation over time during ThermoMaze behavior (Pre, Cooling, Post) and post homecage sleep. Note, that the temperature of the environment was reduced to 10 °C during cooling (yellow line). Brain state classification is shown above the temperature curves (awake, NREM, and REM; black, blue, and red lines, respectively)^44^. **C)** Probability mass function of brain temperature distributions across 10 recording sessions in 2 mice. Cooling and room temperature sub sessions are shown in blue and orange, respectively. **D)** Median brain temperature during cooling and no cooling (room temperature) sessions (not significant, Kolmogorov-Smirnov test). **E)** There is no correlation between brain temperature fluctuation and environmental temperature (linear regression, R = 0.03, p = 0.384; see also Petersen et. al. 2022).

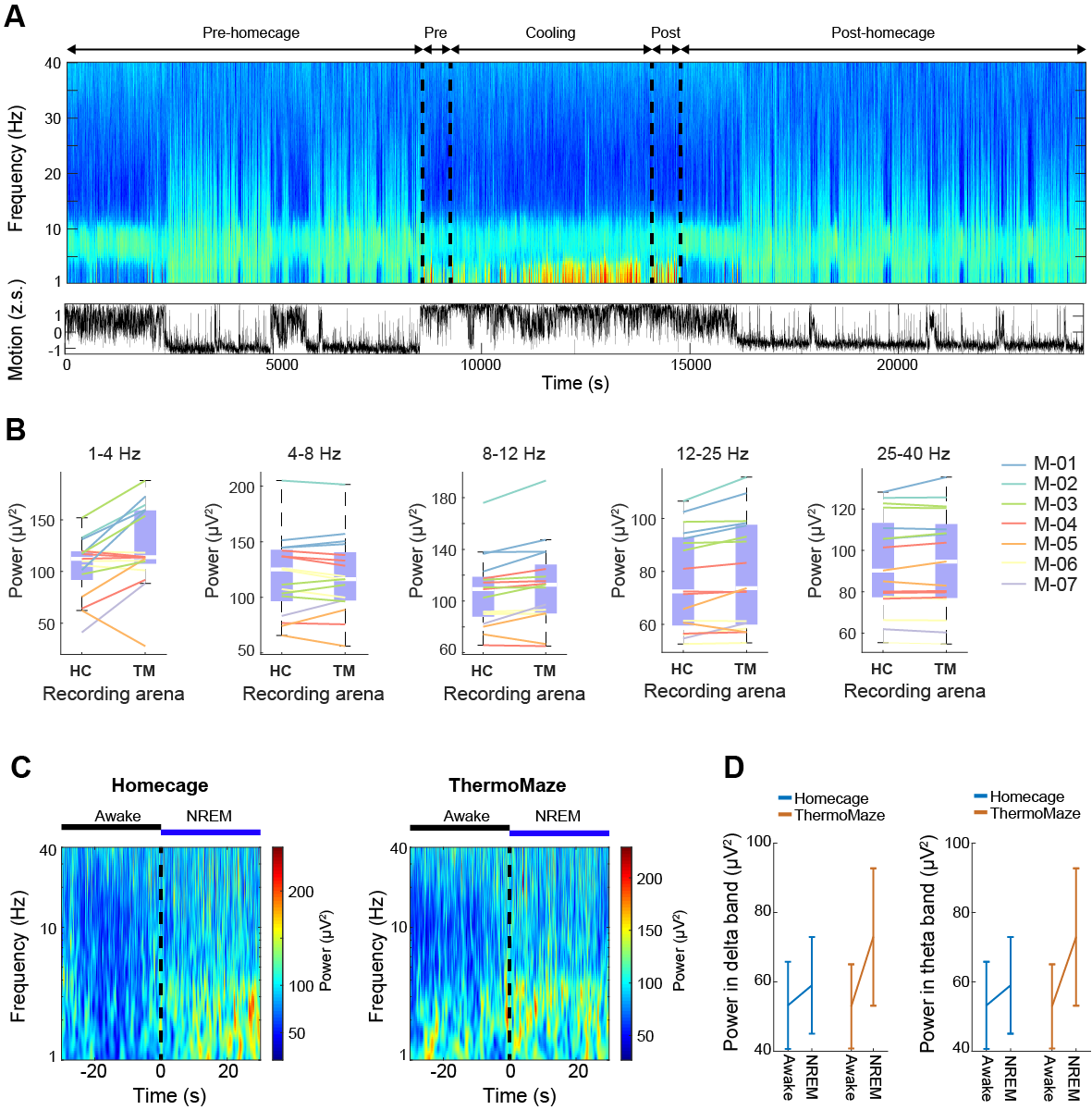

**Supplementary Figure 4. Behavior in the ThermoMaze did not alter hippocampal power spectra. A)** Time-power analysis of hippocampal local field potentials (LFPs). LFP from the CA1 region of the hippocampus was used to calculate the time-resolved fast Fourier transform-based power spectrum (one recording site of a 64-channel silicon probe). Bottom: z-scored motion estimate based on electromyogram (EMG) activity extracted from the intracranially recorded signals (Schomburg, 2014). **B)** Power spectra of the hippocampal LFP (1-40 Hz) were not altered in the ThermoMaze (TM) compared to homecage (HC) during wakefulness (2500 seconds in homecage and ThermoMaze, n = 17 sessions in 7 mice, p > 0.05, Wilcoxon rank sum test). **C)** Awake-NREM transitions (±30 seconds around the transition) triggered power spectrum (n = 7 and 8 transitions in homecage and ThermoMaze, respectively). **D)** Both delta (1-4 Hz) and theta (4-8 Hz) powers were higher in the ThermoMaze following Awake-NREM transitions (n = 7 sessions in 4 mice, p > 0.05, ANOVA paired test).

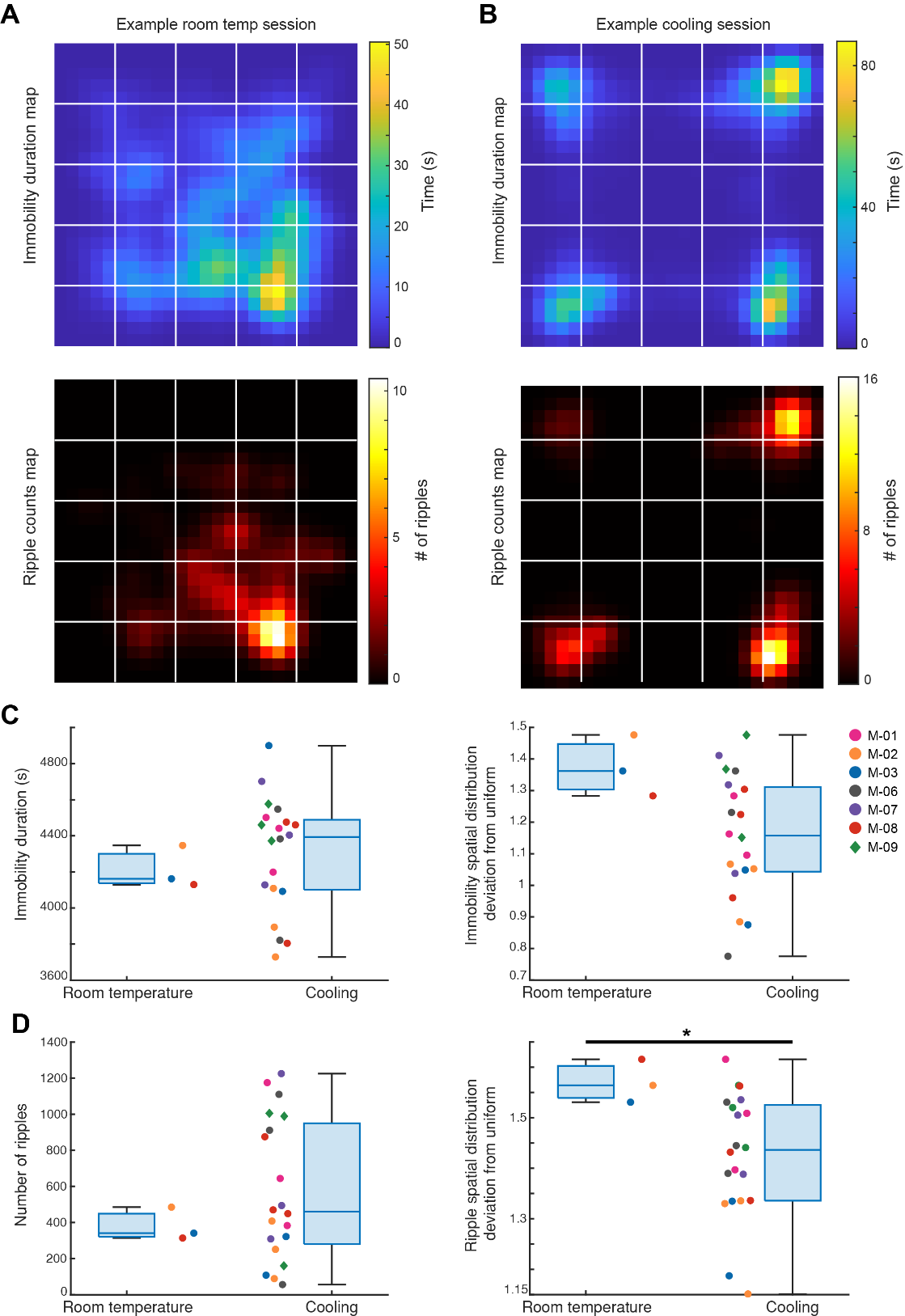

**Supplementary Figure 5. Spatial distributions of immobility duration and SPW-Rr occurrence are more uniform during Cooling compared to room temperature. A)** Top: Immobility duration map of an example session in which the animal was in the ThermoMaze under 25℃ room temperature condition (Mouse_07; Immobility spatial distribution deviation from uniform score: 1.36); Bottom: SPW-R counts map of the same session (SPW-R spatial distribution deviation from uniform score: 1.53). The lower spatial distribution deviation from uniform score indicates that the variable (duration/counts) is more uniformly distributed in the ThermoMaze. **B)** An example Cooling subsession (same plots as in **A**, Mouse_09; Immobility spatial distribution deviation from uniform score: 1.22; SPW-R spatial distribution deviation from uniform score: 1.43). **C)** Left: Immobility durations within an 80-minute period of free exploration of the ThermoMaze either under room temperature or during the Cooling subsession in two groups of mice (room temperature n = 3; Cooling n = 20; p = 0.49). Right: Deviation of spatial distributions of immobility epochs from a uniform distribution (p = 0.08). **D)** Same plots as in **C)** but for total SPW-R counts and the degree to which their spatial distributions deviates from uniform distribution. (Left: p = 0.62; Right: p = 0.04, One-sided Wilcoxon rank sum tests).

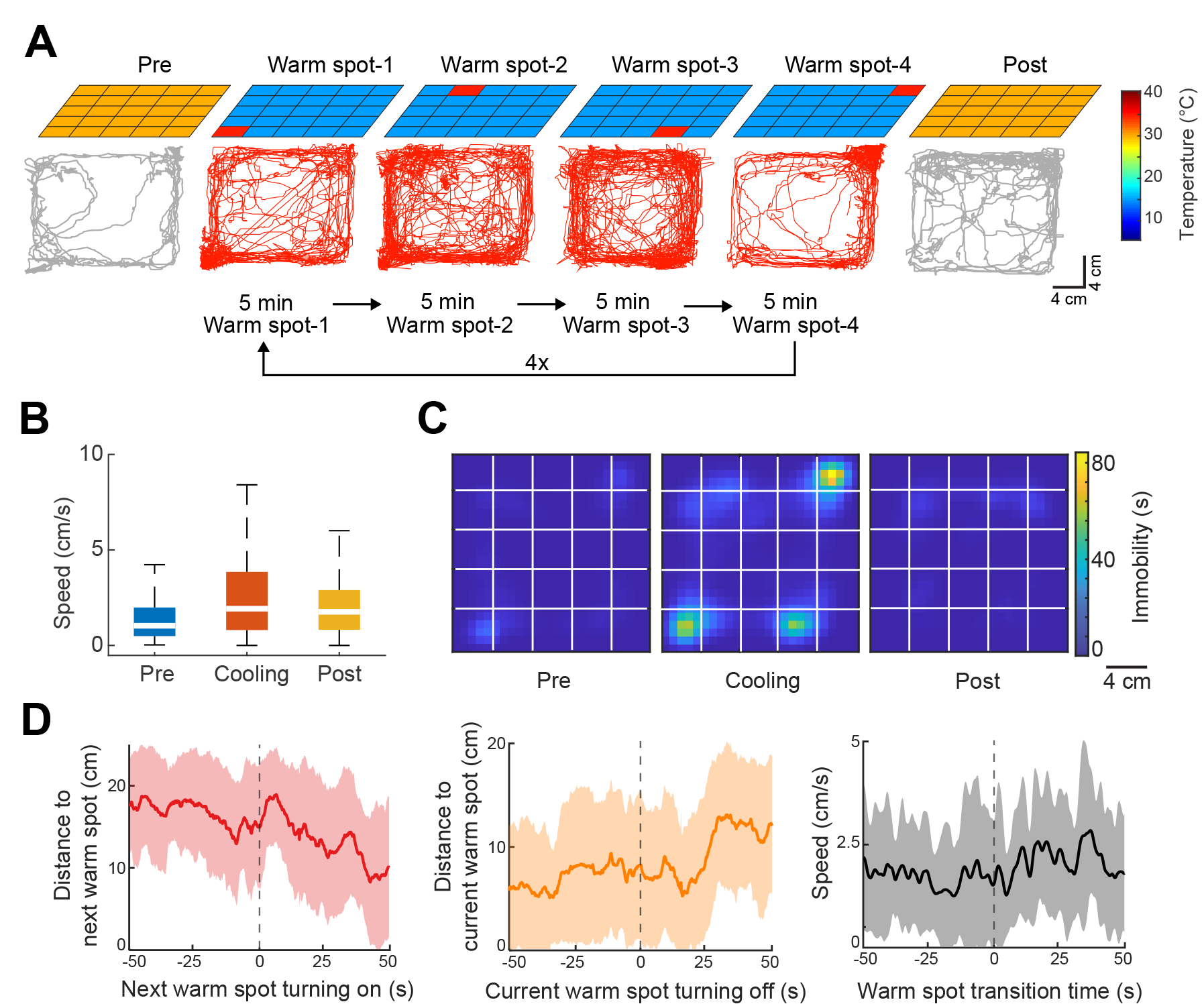

**Supplementary Figure 6. Changing the location of warm spots shape behavior. A)** During Cooling, one of the Peltier elements provided a warm spot for the animal (four Peltier elements, 2 in the corners and 2 close to the corners were used). Each Peltier element was turned on for 5 minutes in a sequential order (1-2-3-4, n = 4 trials). **B)** Animal speed in the ThermoMaze during Pre-cooling (Pre), Cooling and Post-cooling (Post) sub-sessions (n = 3 sessions from n = 2 mice). **C)** Session-averaged duration of immobility (n = 3 sessions in n = 2 mice, speed ≤ 2.5 cm/s) that the animal spent at each location in the ThermoMaze (x and y: animal location (20 x 20 cm); color: temporal duration of immobility (s); white lines represent boundaries of individual Peltier elements). **D)** Left, Median (curve) and 1^st^ to 3^rd^ quartile (shaded region) across sessions of distance to the next warm spot (left panel), and distance from the previous hotspot (middle panel). Right, Speed across sessions centered upon warm spot transition times (time 0).

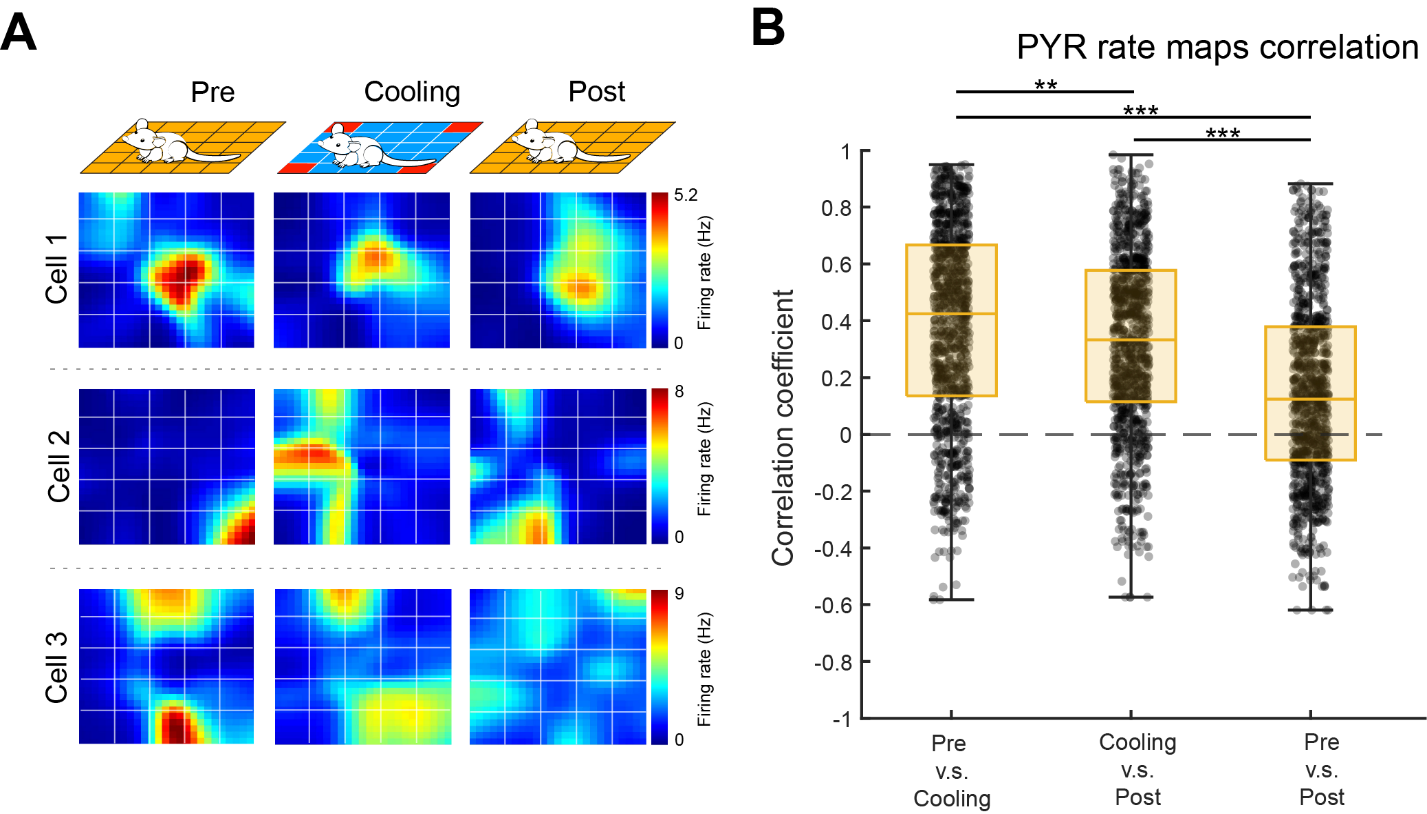

**Supplementary Figure 7. Spatial tuning of hippocampal pyramidal cells in the ThermoMaze. A)** Spatial firing rate maps of three example pyramidal neurons constructed in the three sub-sessions: Pre-cooling (Pre), Cooling and Post-cooling (Post). X and Y: ThermoMaze dimensions; color: firing rate in Hz (color scale is the same across conditions for each cell). **B)** Boxplots of Pearson correlation coefficients between spatial firing rate maps constructed in Pre, Cooling, Post. Median, Kruskal–Wallis test: H = 307.8880, d.f. = 3, p = 0 (n = 1150 pyramidal cells from 7 mice).

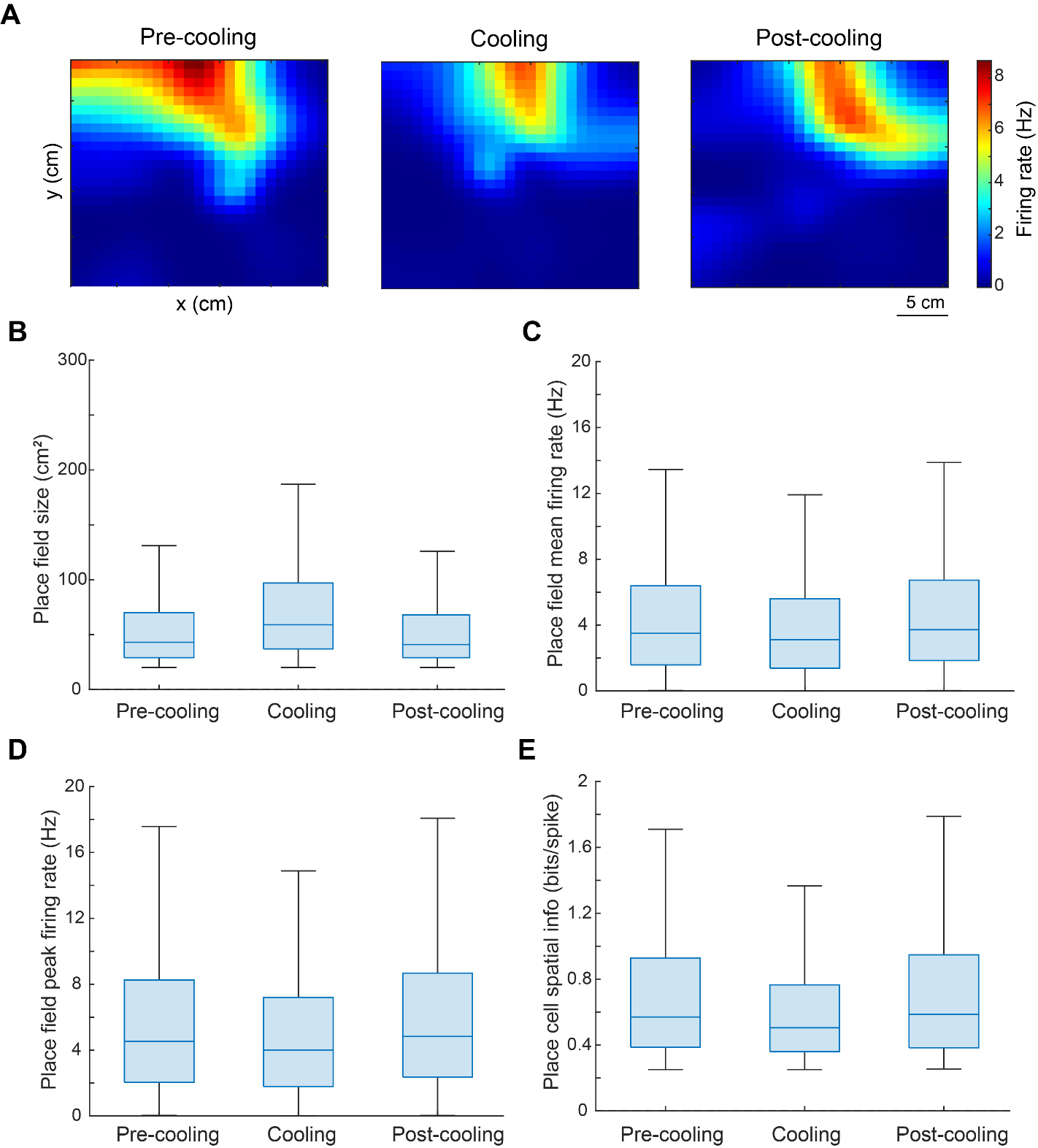

**Supplementary Figure 8. Quantification of spatial tuning properties of CA1 pyramidal neurons as the animal moved through the ThermoMaze. A)** An example neuron showing stable place fields within and across the three sub-sessions. **B, C and D)** Box plot of size (cm^2^), mean firing rate (Hz), and peak firing rate (Hz) of the identified place fields within all individual sub-sessions. The box plot shows the median (center blue line), the lower and upper quartiles (upper and lower blue lines), and the minimum and maximum values that are not outliers (upper and lower black lines). **E)** Box plot of spatial information (in bits/spike) of place cells identified within all sub-sessions.

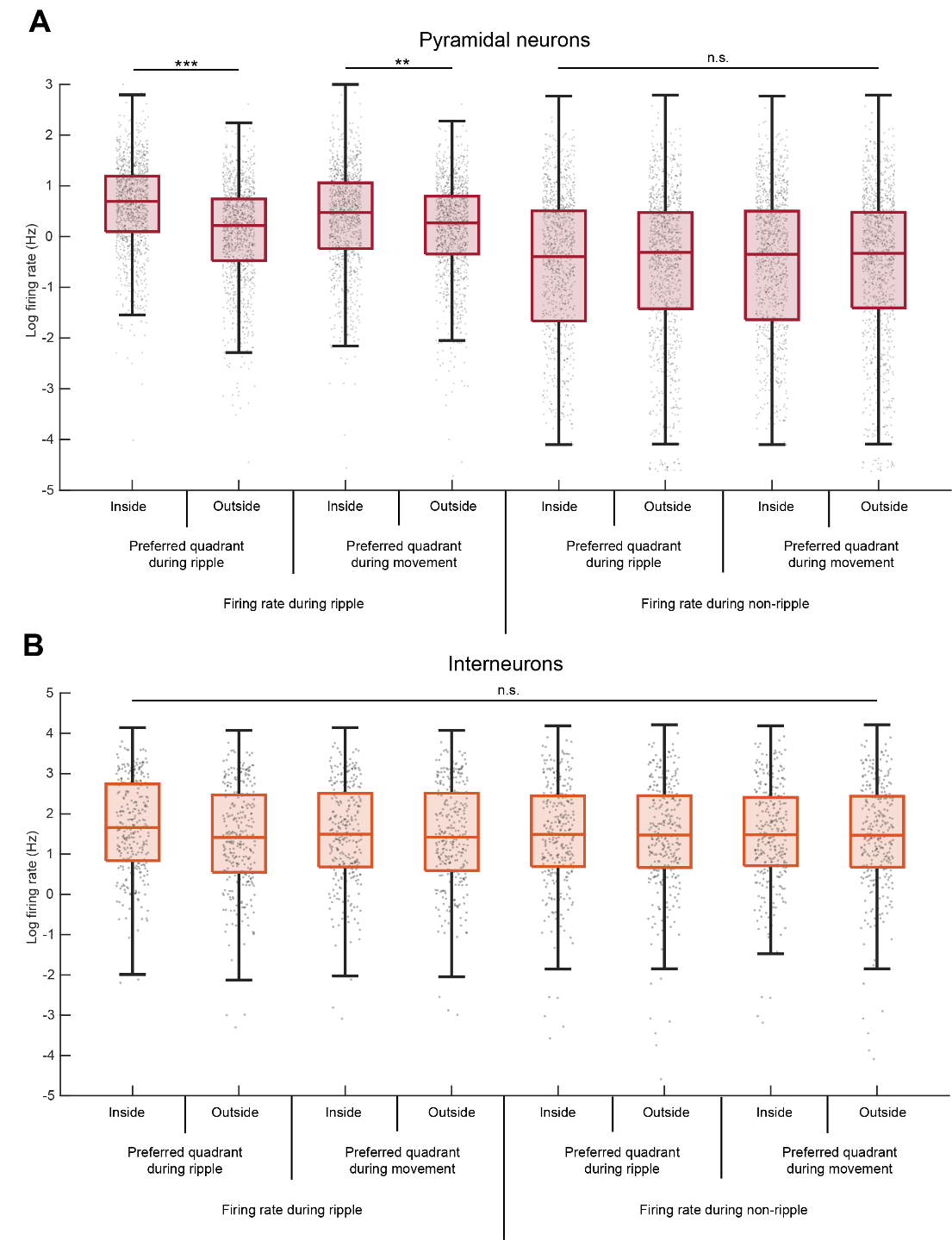

**Supplementary Fig. 9. Comparison of firing patterns of pyramidal cells and interneurons during SPW-Rs. A)** Pyramidal neurons increase their firing rates during SPW-R and movement in their preferred quadrant. Median, Kruskal–Wallis test: H = 992.8856, d.f. = 7, p = 4.1*10^-210^. During SPW-Rs, pyramidal neuron firing rate is significantly higher inside their preferred quadrant (median firing rate = 1.99 Hz) than outside (median = 1.24 Hz), as expected from our definition. This increase in firing rate during SPW-R is also observed when conditioned on inside (median = 1.61 Hz) or outside (median = 1.31 Hz) the cell’s preferred quadrant defined during movement. Firing rate during SPW-R is significantly higher than that during non-ripple (asterisk is omitted in the figure for simplicity; median = 0.67, 0.73, 0.70, and 0.71 Hz for firing rate inside or outside preferred quadrant during ripple or movement, respectively). No significant difference in median is observed among the four conditions for firing rate during non-ripple. **B**) Same as panel **A** but for interneurons. From left to right, median firing rate is 5.26, 4.11, 4.46, 4.14, 4.43, 4.36, 4.39, and 4.34 Hz, respectively. Median, Kruskal–Wallis test: H = 7.1594, d.f. = 7, p = 0.41.

**Supplementary Video 1. Real and thermal image of a mouse in the ThermoMaze.** The animal’s behavior was recorded with a Basler camera and an infrared thermal camera placed above the ThermoMaze. Four Peltier elements were subsequently heated (one in each corner). Infrared image is overlaid on the raw video. The second half of the video is 10 times faster than real time (10 x speed legend in the video).

**Supplementary Video 2. Thermal image of a mouse in the ThermoMaze.** The animal’s behavior was recorded with an infrared thermal camera placed above the ThermoMaze (thermal image is in greyscale). In this video a Peltier element in the inner part of the floor was heated. The speed of the video is 10 times faster than real time.

**Supplementary Table 1. Summary of animal subjects with brain implants**

| **Animal** | **Recording implant** | **Other implants** | **Behavioral protocol** | **# session** | **Sex** |
| --- | --- | --- | --- | --- | --- |
| Mouse_01 | Diagnostic Biochips, 64-2 | NA | ThermoMaze, 4 corners | 1 | F |
| Mouse_02 | Diagnostic Biochips, 64-2 | NA | ThermoMaze, 4 corners | 3 | F |
| Mouse_03 | NeuroNexus, A5x12-16-Buz-Lin-5mm-100-200-160-177 | NA | ThermoMaze, 4 corners | 1 | F |
| Mouse_04 | Tungsten wire | Thermistor | ThermoMaze, 4 corners | 5 | F |
| Mouse_05 | NA | Thermistor | ThermoMaze, 4 corners | 4 | M |
| Mouse_06 | Diagnostic Biochips, 64-2 | NA | ThermoMaze, 4 corners  Inner spots | 3  2 | F |
| Mouse_07 | NeuroNexus A1x32-Poly3-10mm-25s-177 | NA | ThermoMaze, 4 corners  Inner spots | 3  1 | F |
| Mouse_08 | Cambridge Neurotech 64-ch, F6 | NA | ThermoMaze, 4 corners | 2 | F |
| Mouse_09 | NeuroNexus A1x32-Poly3-10mm-25s-177 | NA | ThermoMaze, 4 corners | 4 | M |
| Mouse_10 | NeuroNexus A1x32-Poly3-10mm-25s-177 | NA | ThermoMaze, sleep | 2 | M |
| Mouse_11 | NeuroNexus A1x32-Poly3-10mm-25s-177 | NA | ThermoMaze, sleep | 1 | F |
| Mouse_12 | NeuroNexus A1x32-Poly3-10mm-25s-177 | NA | ThermoMaze, sleep | 2 | M |
| Mouse_13 | Neuropixels 2.0 | NA | ThermoMaze, sleep | 2 | F |

**Supplementary Table 2.** P-values of multiple group comparisons pertaining to analyses of variance in the main figures. Values associated with each group are either means or medians, depending on the statistical test (see main figure legends).

Figure 3C Cumulative distribution of animal speed in the ThermoMaze during three subsessions

| Group A | Group B | p-value |
| --- | --- | --- |
| Pre speed | Cool speed | <0.001 |
| Pre speed | Post speed | <0.001 |
| Cool speed | Post speed | <0.001 |

Figure 4. Boxplots of Pearson correlation coefficients between spatial firing rate maps

Here, group number 1, 2, 3, and 4 refer to correlation values between Pre and Cooling, Cooing and Post and Pre and Post in control sessions.

| Group A | Group B | p-value |
| --- | --- | --- |
| 1 | 2 | 0.006 |
| 1 | 3 | <0.001 |
| 1 | 4 | <0.001 |
| 2 | 3 | <0.001 |
| 2 | 4 | <0.001 |
| 3 | 4 | <0.001 |

Supplementary Figure 7A. Pyramidal neurons increase firing rate during ripples in their preferred quadrant during movement.

Number 1 through 8 represent pyramidal firing rate:

1: during SPW-R inside the cell’s preferred quadrant during ripple

2. during SPW-R outside the cell’s preferred quadrant during ripple

3: during SPW-R inside the cell’s preferred quadrant during movement

4. during SPW-R outside the cell’s preferred quadrant during movement

5: during SPW-R inside the cell’s preferred quadrant during ripple

6. during SPW-R outside the cell’s preferred quadrant during ripple

7: during SPW-R inside the cell’s preferred quadrant during movement

8. during SPW-R outside the cell’s preferred quadrant during movement

| Group A | Group B | p-value |
| --- | --- | --- |
| 1 | 2 | <0.001 |
| 1 | 3 | <0.001 |
| 1 | 4 | <0.001 |
| 1 | 5 | <0.001 |
| 1 | 6 | <0.001 |
| 1 | 7 | <0.001 |
| 1 | 8 | <0.001 |
| 2 | 3 | <0.001 |
| 2 | 4 | 0.596 |
| 2 | 5 | <0.001 |
| 2 | 6 | <0.001 |
| 2 | 7 | <0.001 |
| 2 | 8 | <0.001 |
| 3 | 4 | 0.003 |
| 3 | 5 | <0.001 |
| 3 | 6 | <0.001 |
| 3 | 7 | <0.001 |
| 3 | 8 | <0.001 |
| 4 | 5 | <0.001 |
| 4 | 6 | <0.001 |
| 4 | 7 | <0.001 |
| 4 | 8 | <0.001 |
| 5 | 6 | 0.969 |
| 5 | 7 | 0.9999966894 |
| 5 | 8 | 0.9822369191 |
| 6 | 7 | 0.99416517 |
| 6 | 8 | 0.9999999967 |
| 7 | 8 | 0.9973847873 |

Supplementary Figure 7B. Interneurons firing rate does not change during ripples in their preferred quadrant during movement.

Number 1 through 8 represent interneuron firing rate:

1: during ripples inside the cell’s preferred quadrant during ripple

2. during ripples outside the cell’s preferred quadrant during ripple

3: during ripples inside the cell’s preferred quadrant during movement

4. during non-ripples outside the cell’s preferred quadrant during movement

5: during non-ripples inside the cell’s preferred quadrant during ripple

6. during non-ripples outside the cell’s preferred quadrant during ripple

7: during non-ripples inside the cell’s preferred quadrant during movement

8. during non-ripples outside the cell’s preferred quadrant during movement

| Group A | Group B | p-value |
| --- | --- | --- |
| 1 | 2 | 0.213 |
| 1 | 3 | 0.76q |
| 1 | 4 | 0.454 |
| 1 | 5 | 0.658 |
| 1 | 6 | 0.663 |
| 1 | 7 | 0.604 |
| 1 | 8 | 0.694 |
| 2 | 3 | 0.988 |
| 2 | 4 | 1.000 |
| 2 | 5 | 0.997 |
| 2 | 6 | 0.996 |
| 2 | 7 | 0.998 |
| 2 | 8 | 0.995 |
| 3 | 4 | 1.000 |
| 3 | 5 | 1.000 |
| 3 | 6 | 1.000 |
| 3 | 7 | 1.000 |
| 3 | 8 | 1.000 |
| 4 | 5 | 1.000 |
| 4 | 6 | 1.000 |
| 4 | 7 | 1.000 |
| 4 | 8 | 1.000 |
| 5 | 6 | 1.000 |
| 5 | 7 | 1.000 |
| 5 | 8 | 1.000 |
| 6 | 7 | 1.000 |
| 6 | 8 | 1.000 |
| 7 | 8 | 1.000 |
